# Loss of function, missense, and intronic variants in *NOTCH1* confer different risks for left ventricular outflow tract obstructive heart defects in two European cohorts

**DOI:** 10.1101/215301

**Authors:** Emmi Helle, Aldo Córdova-Palomera, Tiina Ojala, Priyanka Saha, Praneetha Potiny, Stefan Gustafsson, Erik Ingelsson, Michael Bamshad, Deborah Nickerson, Jessica X. Chong, University of Washington Center for Mendelian Genomics, Euan Ashley, James R Priest

**Affiliations:** Children’s Hospital, University of Helsinki, Helsinki, Finland; Cardiovascular Medicine, Stanford University School of Medicine, Stanford, CA; Department of Pediatrics, Division of Pediatric Cardiology Stanford University School of Medicine, Stanford, CA; Department of Medical Sciences, Molecular Epidemiology and Science for Life Laboratory, Uppsala University, Uppsala, Sweden.; Department of Pediatrics, University of Washington, Seattle, WA; Department of Genome Sciences, University of Washington, Seattle, WA; Division of Genetic Medicine, Seattle Children’s Hospital, Seattle, WA

## Abstract

Loss of function variants in *NOTCH1* cause left ventricular outflow tract obstructive defects (LVOTO) in a small percentage of families. Clinical surveys report an increased prevalence of missense variants in *NOTCH1* in family members of individuals with LVOTO and other types of congenital heart disease (CHD). However, the risk conferred by rare variants in *NOTCH1* for LVOTO remains largely uncharacterized. In a cohort of 49 families affected by hypoplastic left heart syndrome, a severe form of LVOTO, we discovered predicted loss of function *NOTCH1* variants in 6% of individuals. Rare missense variants were found in an additional 16% of families. To make a quantitative estimate of the genetic risk posed by variants in *NOTCH1* for LVOTO, we studied associations of 400 coding and non-coding variants in *NOTCH1* in 271 adult cases and 333,571 controls from the UK Biobank. Two rare intronic variants in strong linkage disequilibrium displayed significant association with risk for LVOTO (g.chr9:139427582C>T, Odds Ratio 16.9, p=3.12e-6; g.chr9:139435649C>T, Odds Ratio 19.6, p = 2.44e-6) amongst European-ancestry British individuals. This result was replicated in an independent analysis of 51 cases and 68,901 controls of non-European and mixed ancestry. We conclude that carrying rare predicted loss of function variants or either of two intronic variants in *NOTCH1* confer significant risk for LVOTO. Our approach demonstrates the utility of population-based datasets in quantifying the specific risk of individual variants for disease related phenotypes.

**Author summary:** Congenital heart defects are the most common class of birth defect and are present in 1% of live births. Although CHD cases are often clustered in families, and thus the causal variant(s) are seemingly inherited, finding genetic variants causing these defects has been challenging. With the knowledge that variation in the *NOTCH1* gene previously has been associated with CHDs affecting the left side of the heart, our aim was to further investigate the role of different types of *NOTCH1* variants in left sided CHDs in two cohorts – a cohort of Finnish families with severe lesions affecting the left side of the heart, and the UK Biobank population including individuals with less severe left-sided lesions such as bicuspid aortic valve, congenital aortic stenosis, and coarctation of the aorta. We found a causal loss-of-function *NOTCH1* variant in 6% of the families in the Finnish cohort and in the UK Biobank cohort, we identified two rare variants in the non-coding region of *NOTCH1,* associated with severe left-sided CHDs. These findings support screening of *NOTCH1* loss-of-function variants in patients with severe left sided congenital heart defects and suggests that non-coding region variants in *NOTCH1* play a role in CHDs.

## Introduction

Congenital heart defects (CHDs) are the most common congenital malformations and occur in 0.8–1% of live births [1]. Left ventricular outflow tract obstruction (LVOTO) is a subtype of CHD affecting one or more structures on the left side of the heart – left ventricle, aortic valve and thoracic aorta. At its most severe, LVOTO defects manifest as hypoplastic left heart syndrome (HLHS), in which the left ventricle is underdeveloped, and the systemic circulation depends the persistence of fetal circulatory physiology. Other common LVOTO defects include aortic coarctation (CoA), congenital aortic stenosis (AS), and bicuspid aortic valve (BAV) [1].

The genetic basis of non-syndromic LVOTO defects is largely unknown. Non-syndromic LVOTO defects frequently recur within a family, but often display variable expressivity [2]. LVOTO defects putatively caused by *NOTCH1* variants were initially described in two kindreds with truncating variants [3], and subsequently in several other families [4–12]. In addition to CHDs, *NOTCH1* mutations have been associated with Adams-Oliver syndrome, and certain types of cancers [14,15].

Predicted loss of function (pLOF) variants, e.g. frameshift, nonsense, and splice site variants, in *NOTCH1* have been reproducibly associated with LVOTO defects in multiple studies, and several missense variants have been reported in persons with LVOTO (Fig 1) [3–12]. Previous surveys of LVOTO in both simplex and multiplex families, observed pathogenic or likely pathogenic *NOTCH1* variants in 1–18% of families [5,8,11].

**Figure 1.**
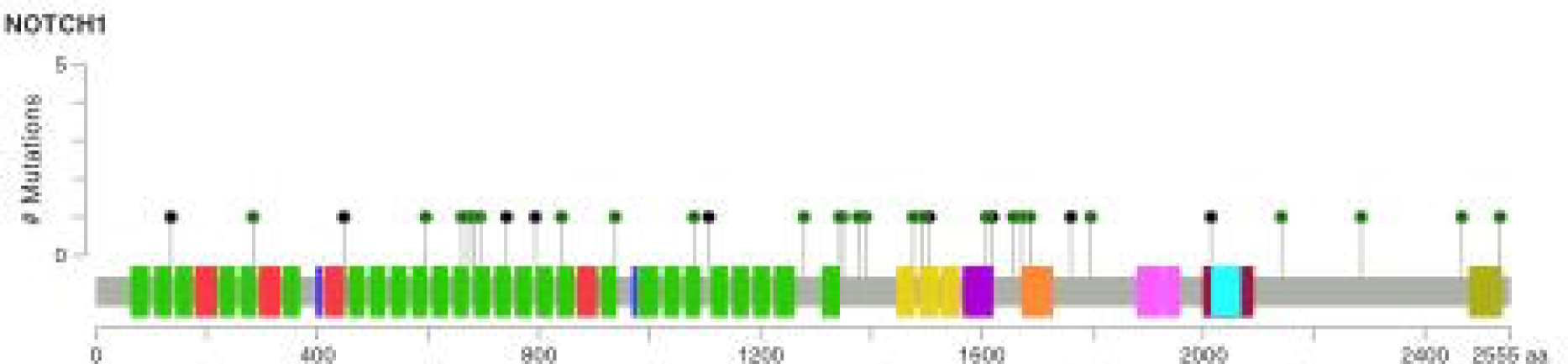
Previously reported non-synonymous and pLOF *NOTCH1* mutations in LVOTO subjects^41,42^.

Both the initial description of *NOTCH1* in LVOTO and subsequent reports include affected members with CHDs other than LVOTO defects including ventricular septal defects and Tetralogy of Fallot (TOF) [3, 11–12]. Additionally, study design has varied among previous analyses rendering estimation of risk and comparisons between studies difficult. Accordingly, while it is clear that pLOF *NOTCH1* variants are associated with LVOTO defects, the role of missense variants and therefore the overall attributable risk of *NOTCH1* variants to LVOTO defects remains unclear.

Here, we describe the presence of pLOF and missense variants in *NOTCH1* in a cohort of 49 simplex cases with HLHS. In addition, we use a large population-based study to assess the risk for LVOTO related heart defects conferred by variants in *NOTCH1* to identify two rare intronic risk variants.

## Results

### Exome Sequencing Reveals Likely Pathogenic Variants and Variants of Unclear Significance

A total of 11 of the 49 probands (22%) had seven protein-altering *NOTCH1* variants, each with a MAF of < 0.05. Three pLOF variants met criteria for pathogenicity (Table 1 and Fig 3). A novel (i.e., absent from all databases) *de novo* truncating variant c.1077C>A (p.Cys359*) with a CADD score of 37 was found in a single HLHS proband. A truncating variant c.1650_1651insT (p.Tyr550*), with a CADD score of 35, was found in a proband with HLHS and CoA and her unaffected parent. p.Tyr550* had been reported previously in a family with Adams-Oliver syndrome (AOS) [20]. A novel splicing variant, c.431-1G>A, with a CADD score of 26 was found in a simplex family with HLHS. This variant was inherited from their father from whom a DNA sample was unavailable.

**Table 1.** Non-synonymous *NOTCH1* variants identified in 49 probands with HLHS

| Diagnosis | Chr | Position | Ref | Alt | dbSNP | DNA change | Protein change | Effect | Allele frequency in gnomAD Finnish | CADD score | Inherited |
| --- | --- | --- | --- | --- | --- | --- | --- | --- | --- | --- | --- |
| HLHS | 9 | 139413065 | G | T | N/A | 1077C>A | Cys359* | Stopgain SNV | 0 | 38,0 | No |
| HLHS, CoA | 9 | 139410452 | G | GT | N/A | 1650_1651insA | Y550_T551* | Stopgain SNV | 0 | 35,0 | Yes |
| HLHS | 9 | 139404414 | C | T | N/A | c.2741-1G>A |  | Splicing | 0 | 26,0 | Yes |
| 5 HLHS, 1 BAV* | 9 | 139401233 | C | T | rs61751543 | 3836G>A | Arg1279His | ns SNV | 0,02097 | 16,2 | Yes 2/ Not known 3 |
| HLHS | 9 | 139401216 | C | T | N/A | 3853G>A | Val1285Met | ns SNV | 0 | 29,8 | 29,8 |
| HLHS | 9 | 139391338 | C | T | rs61751489 | 6853G>A | Val2285Ile | ns SNV | 0,00617 | 0.5 | Not known |
| HLHS, BAV, CoA, HAA, ASD, VSD | 9 | 139391013 | T | C | N/A | 7178A>G | Gln2393Arg | ns SNV | 0 | 10,1 | Not known |

\* 6 affected

Among the four remaining *NOTCH1* variants, a missense variant c.3836G>A (p.Arg1279His) was found in 5 HLHS probands and in one person with BAV who was a half-sister of a proband. p.Arg1279His has been found in both persons with LVOTO and in controls in three previous studies [6,7,9]. Another missense variant, c.6853G>A (p.Val2285Ile), was found in one child with HLHS and his unaffected parent. A novel missense variant, c.7178A>G (p.Gln2393Arg), was found in a proband with HLHS, BAV, CoA, HAA, ASD, VSD. Finally, a rare missense variant c. 3853G>A (p.Val1285Met), was found in a singleton proband with HLHS.

### Rare intronic variants in *NOTCH1* are associated with risk for LVOTO in European and non-European populations

Given the uncertain of whether the four missense variants detected in the probands with HLHS were causal, we decided to test the association of common and rare coding and noncoding variants in *NOTCH1* with risk of LVOTO and related CHD phenotypes more broadly in UK Biobank, a large population-based study. We first developed a classification scheme to identify 396 cases with LVOTO in a highly specific manner. To determine the power for detecting associations with rare-variants, we performed simulations of the Firth’s penalized regression. As BAV was included in our definition of LVOTO and may commonly remain undiagnosed in the population at a significant rate, our simulations included misspecification of cases as controls. Power was largely dependent on the minor allele frequency, with misspecification of cases playing a negligible role in power at all simulated rates of misspecification of cases (S1 Fig). For the combined hybrid-LVOTO phenotype (n= 271) the power to detect a genetic association with even low-risk variants of OR 2 or greater was nearly 100% at a minor allele frequency of 0.001, and at a minor allele frequency of 0.0001 nearly 80% to detect variants conferring a risk of 12.9 or higher (S1 Fig). These simulations provide evidence of adequate power to detect genetic associations for CHD phenotypes related to *NOTCH1* even in the presence of unrecognized cases within a large control population.

We identified 400 non-coding or coding (missense and synonymous) variants in *NOTCH1* and no pLOF variants (S1 Table) available for analysis imputed with high quality. Among these 400 variants, none of the coding variants met our pre-specified threshold for locus wide significance. Two of the four non-synonymous variants we observed Finnish cohort (Val2285Ile and Arg1279His) were present in the UK biobank cohort but were not significantly associated with CHD.

Associations the hybrid LVOTO phenotype with two rare intronic variants in *NOTCH1* (g.chr9:139427582C>T, Odds Ratio 16.9, p=3.12e-6 and g.chr9:139435649C>T, Odds Ratio 19.6, p = 2.44e-6) met the locus-wide threshold for significance (Fig 4). As these two intronic variants appear at similar frequencies within non-European populations in the gnomAD database of population level genomic variation, we repeated analysis limited to the 68,952 individuals of non-European and mixed ancestry within the UK Biobank. The association with risk for LVOTO identified was of the same direction and similar magnitude (g.chr9:139427582C>T, Odds Ratio 12.3, p=0.019 and g.chr9:139435649C>T, Odds Ratio 13.9, p = 0.014) (Fig 4 and Fig 5).

**Figure 2.**
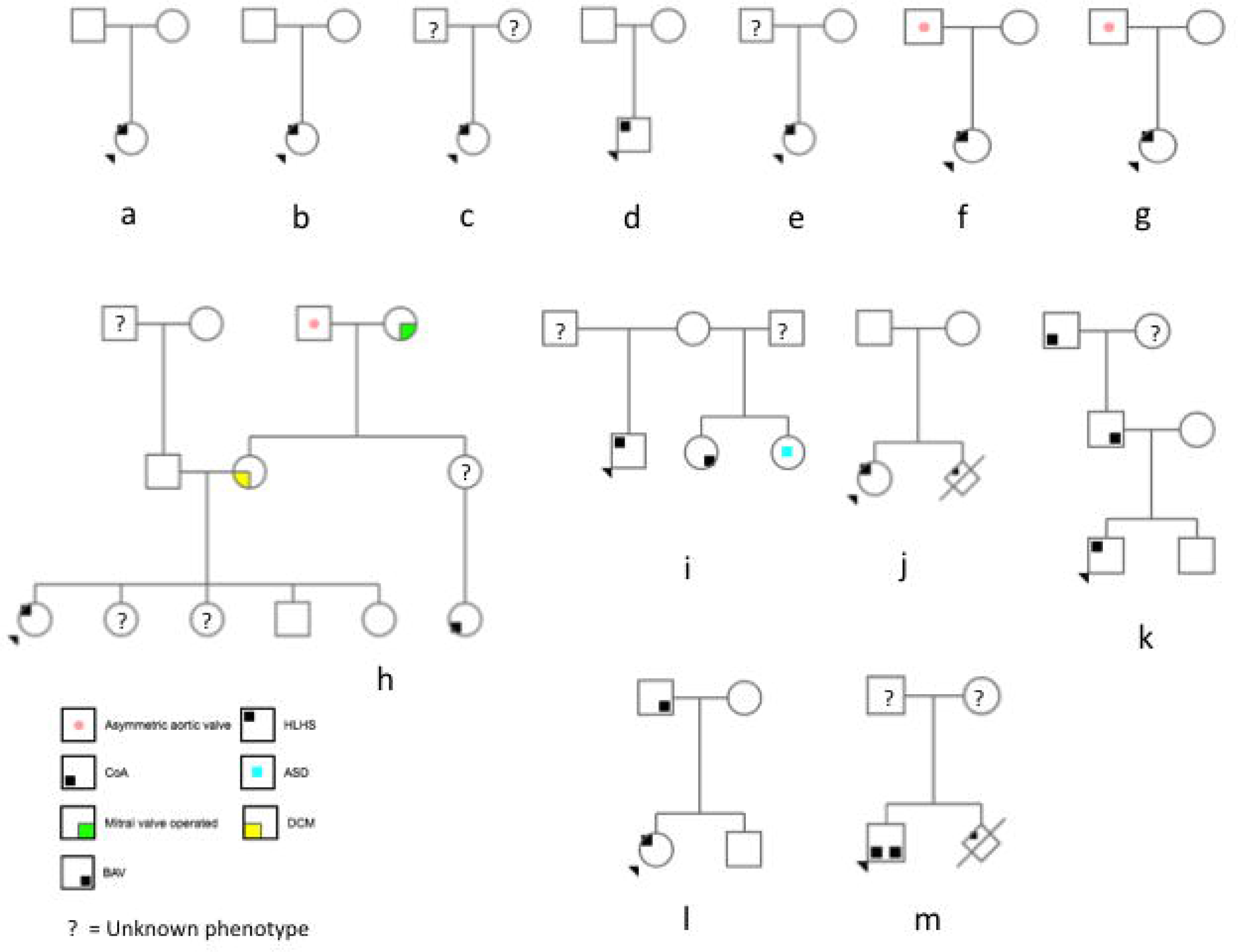
Pedigrees of 5 trios with unaffected (a-c), unknown (d, e), and possibly affected (f, g) parental phenotypes, and pedigrees of 6 families with more than one affected member (h-m).

**Figure 3.**
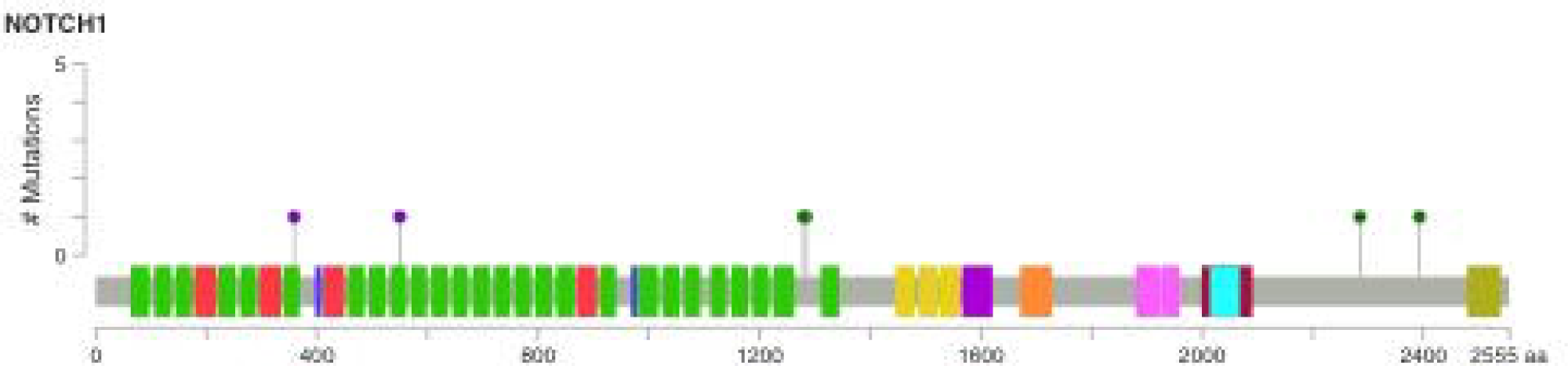
Non-synonymous and pLOF mutations in NOTCH1 in the study population presented in Mutation Mapper as a lollipop plot^41,42^.

**Figure 4.**
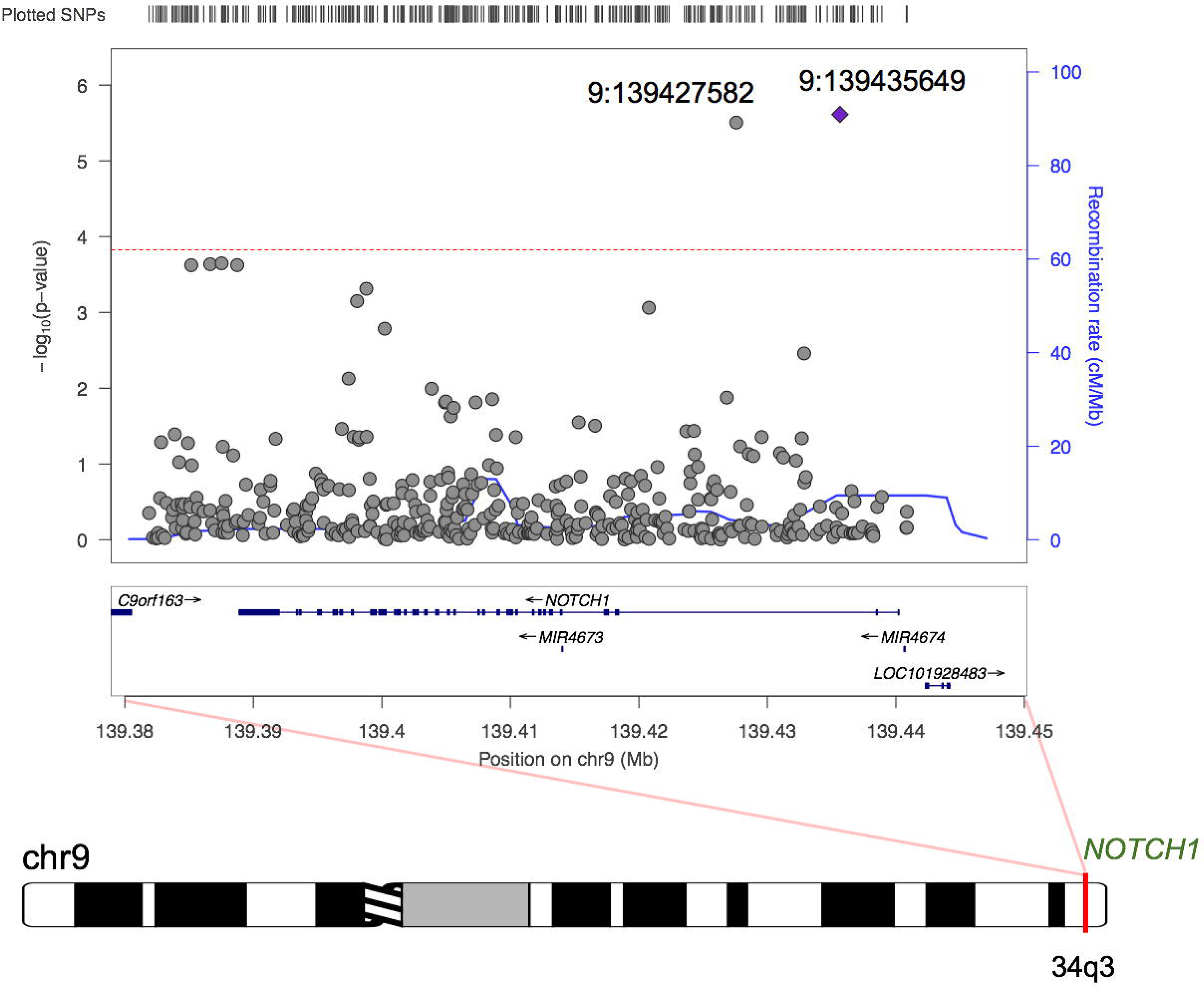
Region plot for the association of *NOTCH1* variants with LVOTO.

**Figure 5.**
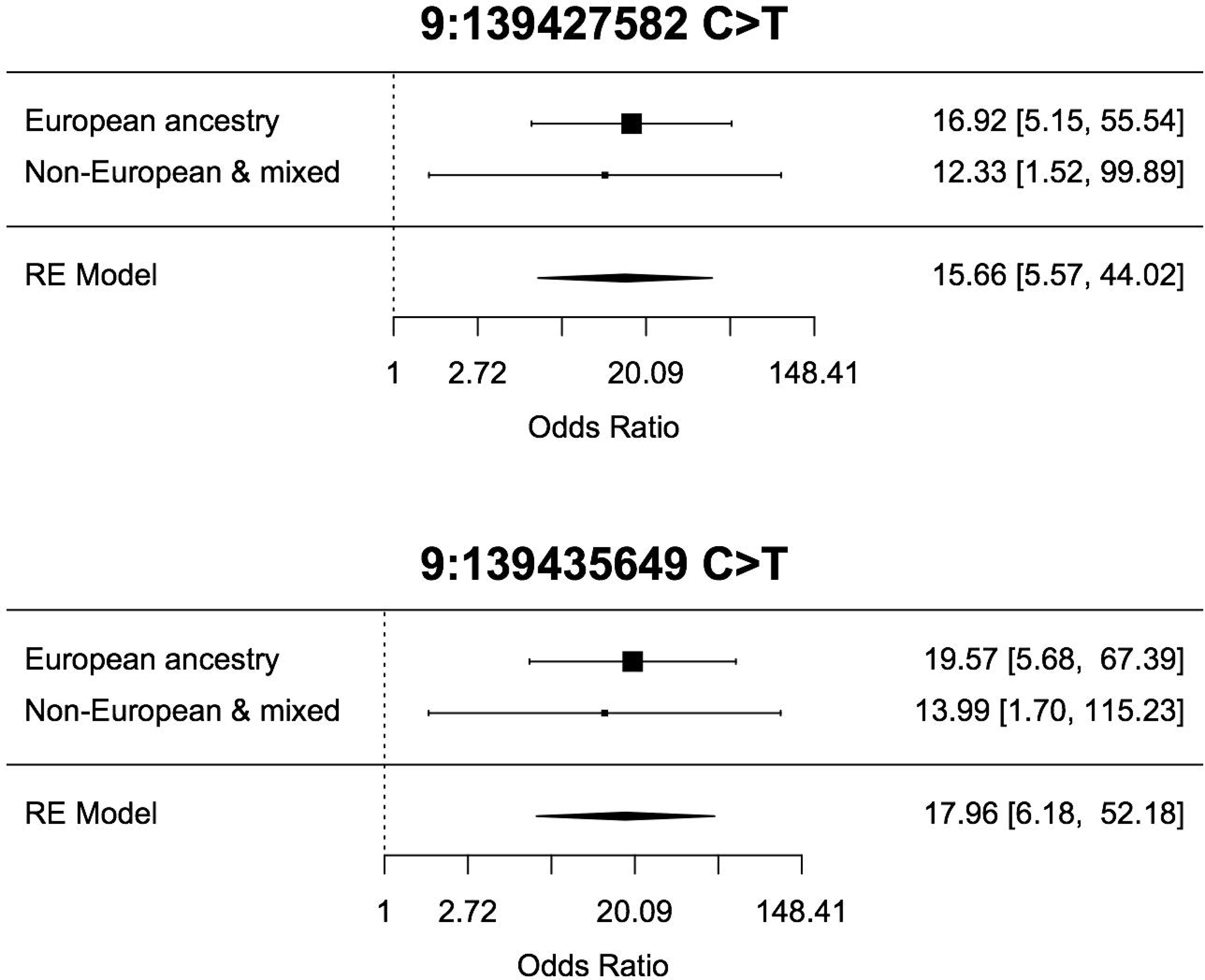
Effect sizes of the associated *NOTCH1* variants in European and non-European and mixed populations.

## Discussion

We detected a likely pathogenic/pLOF mutation in *NOTCH1* in 6% of individuals with HLHS in a Finnish cohort [21]. These variants included a splicing variant and two truncating variants. These findings suggest that pLOF variants in *NOTCH1* may be sufficiently prevalent in LVOTO defects to warrant genetic testing. An additional 16% of HLHS probands in this cohort had missense variants of unknown significance. Further study of missense *NOTCH1* variants in a large population-based study of LVOTO defects did not reveal any significant association between missense variants and risk for less severe LVOTO-related defects. In the population-based study, neither common nor rare missense variants in *NOTCH1* were significantly associated with LVOTO defects. Notably, no pLOF variants were identified in the population-based study cohort, suggesting that such variants are very rare in putatively healthy population controls.

Two rare intronic variants displayed strong association with risk for LVOTO defects in both European and non-European/mixed populations. These two variants appear to exist in similar frequencies in European (MAF=0.00074), Ashkenazi Jewish (MAF=0.0078), and African (MAF=0.0003) populations, and are separated by 8,068 base pairs and display strong linkage disequilibrium (*r*^2^ = 0.949) suggesting the associations with LVOTO are not unique or independent. The two variants exist within the large intron between the second and third exons of *NOTCH1*, and are not located with sufficient proximity to exert a functional effect on canonical splicing of the transcripts for *NOTCH1* or nearby microRNAs (*MIR4673* and *MIR4674*) and additionally do not appear to be strongly conserved through evolution. Thus a direct effect of the two variants upon the protein sequence or structure of NOTCH1 is not clear and will require an experimental approach to determine function which may be related to transcription or epigenetic regulation.

The absence of a significant association of missense variants in *NOTCH1* with LVOTO-related phenotypes in UK Biobank must be interpreted with attention to the characteristics of the study population. Long-term survival of patients with HLHS was achieved by surgical innovations during 1980’s, and thus there were no patients with HLHS within the UK Biobank cohort, which consists of middle-aged to elderly individuals from the general population. The small fraction of individuals classified with LVOTO is lower than previous population estimates of BAV (the most common type of LVOTO) suggesting some ascertainment errors due to the use of phenotyping from medical records, perhaps in a combination with a healthy cohort effect (the individuals in UK Biobank being healthier than the general population).

Moreover, as pathogenic *NOTCH1* mutations have been found more frequently in pediatric study cohorts than in adult cohorts, it has been speculated that *NOTCH1* mutations are more often found in severe disease [11]. Cardiac MRI data are currently available for a small fraction of individuals within the UKBiobank. Of the LVOTO cases classified by the specification schema, two individuals identified had available MRI imaging, and both were positively identified as having bicuspid aortic valve. Our algorithmic approach to classification of individuals with CHD relying upon clinical and self-reported data requires additional validation, but the overall approach may offer insight into rare alleles conferring risk for disease that has been obscured by the underlying genetic architecture of complex diseases in diverse human populations [22].

Two of our 49 probands had novel truncating variants, and truncating variants in *NOTCH1* have been reproducibly associated with LVOTO defects. A total of 14 of the 138,632 gnomAD subjects have truncating *NOTCH1* variants and the probability of loss of function variant intolerance (pLI) is estimated to be 1.0. Previously both *de novo* [10] and familial [11] splice site variants have been reported in LVOTO subjects. The splice site variant 431-1G>A found in an HLHS proband in this study was also found in the asymptomatic father indicating reduced penetrance, which is in accordance with a previous study where familial splicing variants had 87% penetrance [11]. According to the Human Splicing Finder, the 431-1G>A variant alters the WT acceptor site and is likely to affect splicing [25]. Only 5 of 138,632 individuals included in gnomAD have splice site variants in *NOTCH1*, indicating these are poorly tolerated variants.

The truncating p.Tyr550* variant found in one of the HLHS probands studied herein has been previously reported in a kindred with four affected members with AOS with variable expression of congenital limb defects and scalp cutis aplasia [20]. Of these four AO individuals, one had undergone cardiac evaluation by echocardiogram showing AS and aortic regurgitation. Functional analysis associated the variant with reduced expression of *NOTCH1, HEY1* and *HES1* in peripheral blood as measured by RT-PCR indicating that the truncated protein is likely to be subject to nonsense-mediated decay reducing the downstream *NOTCH1* signaling. Notably, two family members with this variant did not have AOS; however, one of them had an unexplained heart murmur, and the other had aortic regurgitation, and a family member with unknown genetic variant status died of an unspecified CHD.

Based on our analyses in the UK Biobank, which failed to detect a significant association of missense variants in *NOTCH1* with LVOTO-related phenotypes, we think the four non-synonymous variants are not able to cause disease in isolation. However, the inheritance pattern of CHD is in many cases complex, and the contribution of these variants to disease together with other predisposing environmental of genetic factors cannot be determined. The Arg1279His variant (rs61751543) is particularly interesting, as it has been seen more frequently in cases vs. controls in three previous LVOTO cohorts[6,7,9]. In this study it is seen in 5/49 (10%) HLHS probands compared to 2% of the Finnish gnomAD population. The variant has a CADD score of 16.23, which could be interpreted to suggest potential pathogenicity *in silico*. Notably, the Finnish gnomAD population has not been phenotyped for CHD, and likely contains some sporadic individuals with undiagnosed BAV, which is relatively common. However, recent analyses incorporating variant penetrance and incomplete ascertainment of control populations [26] suggest that a conservative view must be taken when assigning disease risk for individual variants. Additionally, in functional analysis the Arg1279His variant does not diminish *JAG1* induced *NOTCH1* signaling [6]. Functional studies on the role of low-frequency variants overrepresented in CHD in patient-derived induced pluripotent stem cells would be fruitful in assessing their pathogenic potential.

The discovery of the same truncating variant in a family with HLHS and a family with AOS with minor or no detectable cardiac phenotype is illustrative of variable phenotypic expressivity. Different mutations in the same gene causing different phenotypes in different families may be due to the presence of modifying mutations within a network of interacting proteins [27], stochastic differences in transcriptional dynamics or cardiovascular development, poorly characterized aspects of epigenetic inheritance, or environmental factors. Given that epistatic effects between genetic variants are difficult to detect even in large genetic studies [28], discovery of modifying factors (genetic, environmental, epigenetic, or otherwise) may depend on hypothesis-driven experimental approaches [29].

In conclusion, in our study of 49 HLHS individuals, three (6%) had loss of function variants, which are likely causative for the congenital heart defects. In addition, five affected individuals (10%) and one affected relative displayed a low frequency variant p.Arg1279His present in 2% of the general population which does not appear to be associated with risk for disease in a large-scale association study of less severe phenotypes. Two rare intronic variants displayed strong association with risk for LVOTO defects. However, due to their deep intronic location, the direct functional effect of the two variants upon *NOTCH1* is not clear. Our finding of pathogenic *NOTCH1* variants in 6% of the study subjects is similar in prevalence to single genes causing Long QT syndrome and hypertrophic cardiomyopathy, cardiac conditions for which genetic testing is routine. Thus our data are supportive of the use of clinical testing for *NOTCH1* variant screening patients with HLHS and other LVOTO.

Rare, highly penetrant LOF variants clearly increase risk for nonsyndromic CHD[30,31] in a small percentage of the general population while the role of missense variants remains unclear. Genetic risk factors that explain the bulk of syndromic and non-syndromic CHD remain to be discovered. Moreover, the role of environmental modifiers and interactions among loci remain largely unexplored. Access to large cohorts of robustly phenotyped families with CHD and the availability of comparative genomic data sets from large reference populations will enable both careful assessment of the pathogenicity of rare variants and facilitate identification of novel variants / genes associated with CHD. Functional studies of the pathogenic potential of such variants in patient-derived induced pluripotent stem cells may confirm the pathogenicity of these variants and may serve to elucidate mechanisms behind reduced penetrance. For genes where causality for cardiac malformations is well established, our findings may suggest an opportunity to quantify risk conferred by inherited alleles to increase the known fraction of genetic attributable risk for CHD.

## Methods

### Exome Sequencing in Finnish Probands and Relatives

We recruited a cohort of 49 patients with HLHS from Helsinki University Children’s Hospital. Exome sequencing was performed by the University of Washington Center for Mendelian Genomics Seattle, USA, on DNA samples collected from 37 probands, 7 trios (5 with unaffected or unknown parental phenotypes, and 2 with one affected parent), and 5 probands with a family history of LVOTO defects. In these latter cases, we sequenced the parents, siblings and other affected family members (Fig 2a-c). All persons sequenced were of self-reported Finnish ancestry.

In brief, library capture was performed with Roche/Nimblegen SeqCap EZ v2.0, with 75-base pair paired-end sequencing on the HiSeq2500/4000 instrument (RTA 1.18.34/RTA 2.5.2). BAM files were aligned to a human reference (hg19hs37d5) using BWA-MEM (Burrows-Wheeler Aligner; v0.7.10) [32]. Read-pairs not mapping within ± 2 standard deviations of the average library size (~150 ± 15 bp for exomes) were removed. RTG-core version 3.3.2 was applied to the raw exome sequence data for mapping, pedigree-aware variant calling, and genotype filtration (Real Time Genomics Inc., Hamilton, New Zealand) [33].

We analyzed all non-synonymous *NOTCH1* variants found with an allele frequency of <0.05 in the Genome Aggregation Database (gnomAD) [34]. In addition, we checked the occurrence and frequency of the candidate variants in the Sequencing Initiative Suomi (SISu) [35] database. Annotation was performed with the internally developed STANNOVAR tool [36]. Combined Annotation Dependent Depletion (CADD) scores were used to estimate the pathogenicity of the candidate variants. CADD is a tool for scoring the deleteriousness of single nucleotide variants as well as insertion/deletions variants in the human genome. The CADD score integrates multiple annotations into one metric by contrasting variants that survived natural selection with simulated mutations [37]. Likely pathogenic variants were confirmed with Sanger sequencing of PCR amplicons in all available family members. The following PCR primers were used: NOTCH1 p.Tyr550Ter: Forward-GCACACTCGTTGATGTCCTC, Reverse-AGAACTGTCTCTCCTCCCCT; NOTCH1 c.431-1G>A: Forward-TACTCAGGATTGGGGCTGAG, Reverse-GAAGGGCCATAGTGCTGTTG; NOTCH1 p.Cys359Ter: Forward-GTTGTAAAACGACGGCCAGTGTGAGGTCACACAGCTCAGG, Reverse-TCACACAGGAAACAGCTATGAGTACCGAGGATGTGGACGAG.

The guidelines of the Declaration of Helsinki were followed and the study was approved by the Ethics Board of Helsinki and Uusimaa Hospital District. Written informed consent was obtained from each participant over 6 years of age, and from both parents of each minor participant.

## Population Studies in the UK Biobank

### Classification of Left Ventricular Outflow Tract Disease from Biobank Data

A classification algorithm (S2 Fig) was developed for defining case and control subjects using diagnostic codes from the International Classification of Diseases versions 9 and 10, the OPCS Classification of Interventions and Procedures version 4, and from self-reported medical history data collected in a questionnaire and codified by a trained nurse for the UK Biobank study (all codes listed in S3 Table). The LVOTO phenotype was defined as comprising the following individual phenotypes of congenital etiology: aortic stenosis, subaortic stenosis, aortic insufficiency, aortic coarctation, aortic atresia, congenital aneurysm of the aorta, and hypoplastic left heart syndrome. Since an undiagnosed bicuspid aortic valve may manifest with outflow obstruction (i.e. aortic stenosis and/or insufficiency) later in life, the classification algorithm was designed to identify patients with bicuspid aortic valves who otherwise were not codified as having a congenital heart defect, a process not previously defined in the literature. Patients with aortic valve disease with unspecified etiology or who had had an aortic valve replacement were excluded from cases and reclassified as controls if they met criteria for non-congenital etiologies of valve disease (i.e. rheumatic heart disease, endocarditis, etc.) (S3 Table). Furthermore, of the remaining cases, if the age at diagnosis was >45 for aortic valve disease and age at surgery was >50 for valve replacement, then the patient was excluded to reduce the chance of false positives due to age-related degenerative aortic valve disease. False positive rates were predicted using data on the frequency of bicuspid aortic valves by decade of life published in 2011 by Roberts and Ko [38]. Lastly, control subjects meeting criteria for diagnoses that are associated with syndromic or sporadic congenital heart disease (i.e. endocarditis, thoracic aortic aneurysm, aortic root dilation) but otherwise unable to meet inclusion criteria were excluded to minimize the chance of false negatives in the control population (S3 Table).

### Association Testing for *NOTCH1* Variants

We performed a primary genetic association study for LVOTO and related phenotypes in 271 cases and 333,571 controls of European ancestry included in the recent release of imputed genetic data from the UK Biobank [39,40]. From the dataset of imputed variants, we analyzed 400 common and rare variants in *NOTCH1*. Summary statistics from the association tests of European ancestry participants can be found in S4 Table.

Statistical testing was performed by standard methodology using PLINK version 2.0 using the hybrid logistic regression with Firth’s penalized regression fallback for non-converging models with PLINK’s *--glm firth-fallback* option) [41,42]. and included 10 principal components related to ancestry as continuous covariates and two binary covariates related to genotyping batch. From the imputed dataset, we included 400 exonic and intronic variants with 30 or more alleles in the population that had missing rates of 5% or lower and high quality imputation (95% or greater individuals for which the maximum genotype probability was greater than a threshold of 0.9), and which also displayed an empirical-theoretical variance ratio (MaCH’s *r*^2^) >0.8. For analysis of any single variant, individuals with missing calls were excluded. Primary testing was performed in 271 cases and 333,571 controls of white British origin. Confirmatory testing was performed in 51 cases and 68,952 controls of non-European or mixed ancestry.

We performed a single analysis of the LVOTO hybrid, and estimates of risk ratios and confidence intervals for individual variants were obtained including the same covariates, as mentioned above. Significance thresholds were predetermined; we report a locus-wide 0.000125 (0.05 / 400 analyzed variants in *NOTCH1*) significance threshold.

Power calculations were performed for unbalanced case control study design employing the *logistf* package implementation of the Firth’s penalized logistic regression in the R language for statistical computing [42]. Firth’s regression implementation in the *logistf* package is used by PLINK 2 for the association tests when conventional models fail to converge. Simulations were conducted in 1000 replicates holding constant a variant with a minor allele frequency of 0.1, 0.05, 0.01, 0.001, or 0.0001 and a control population size of 333,571 individuals, with variation of the genotype relative risk (GRR) at 2, 4, 8, 16, and 32. For simplicity the simulations employed an additive model of risk, for which the GRR = *f*_*1*_ / *f*_*0*_ where *f*_*0*_ and *f*_*1*_ represent the likelihood of being affected with LVOTO for individuals with 0 or 1 risk alleles respectively. Additionally, as our classification schema for cases was weighted towards specificity (described above) and the high likelihood of misspecification of undiagnosed BAV within the control population, we simulated mis-specification of controls at a rate of 0.01% 0.1%, and 1%. Estimation of power for specific risk and allele frequencies was calculated using local polynomial regression from the simulated datasets.

## Acknowledgements

Sequencing was provided by the University of Washington Center for Mendelian Genomics (UW-CMG) and was funded by the National Human Genome Research Institute and the National Heart, Lung and Blood Institute grant HG006493 to Drs. Deborah Nickerson, Michael Bamshad, and Suzanne Leal.

